# Harbor porpoise losing its edges: genetic time series suggests a rapid population decline in Iberian waters over the last 30 years

**DOI:** 10.1101/2021.08.19.456945

**Authors:** Yacine Ben Chehida, Tjibbe Stelwagen, Jeroen P. A. Hoekendijk, Marisa Ferreira, Catarina Eira, Andreia T. Pereira, Lidia Nicolau, Julie Thumloup, Michael C. Fontaine

## Abstract

Impact of climate changes on species is expected to be especially visible at the extremities of the species distribution, where they meet sub-optimal conditions. In Mauritania and Iberia, two genetically isolated populations of harbor porpoises form a distinct ecotype and are presumably locally adapted to the upwelling waters. By analyzing the evolution of mitochondrial genetic variation in the Iberian population between two temporal cohorts (1990-2002 vs. 2012-2015), we report a dramatic decrease in genetic diversity. Phylogenetic analyses including neighboring populations identified two porpoises in southern Iberia carrying a divergent haplotype close to the Mauritanian population, yet forming a distinctive lineage. This suggests that Iberian porpoises may not be as isolated as previously thought with immigration from Mauritania or an unknown population in between, but none from the northern ecotype. The rapid decline in the Iberian mitochondrial diversity may be driven by either genetic drift, or by a dramatic decline in census population size possibly resulting from environmental stochasticity, prey depletion, or acute fishery bycatches. These results illustrate the value of genetics time series to inform demographic trends and emphasize the urgent need for conservation measures to ensure the viability of this small harbor porpoise population in Iberia.

## 1. Introduction

The impact of climate changes on species is expected to be especially visible at their distribution edges, where species are expected to meet the limits of their ecological tolerance. Marginal populations at the rear (warm) distribution edge are often considered more susceptible to warming than central populations because of the warmer ambient temperatures they experience, but this overlooks the potential for local adaptation (Bennett et al., 2015; Vilà-Cabrera et al., 2019).

Cetaceans may be particularly vulnerable to the current climate change (MacLeod, 2009) despite their crucial function as apex predators in marine ecosystems (Bowen, 1997). Climate change influences the availability of their prey and modifies the habitat, thereby impacting several (if not all) cetacean species and may even put some of them at a high risk of extinction (Learmonth et al., 2006). As a response, several cetacean species are shifting their distribution ranges to track suitable habitats or adapt locally by switching to different food resources (Lambert et al., 2011; Learmonth et al., 2006; MacLeod, 2009; Williamson et al., 2021). For example, temperate and subpolar small cetaceans like harbor porpoises (*Phocoena phocoena*) are likely to show a poleward shift (Heide-Jørgensen et al., 2011; MacLeod, 2009). Due to an elevated metabolic rate and fast reproductive turnover, this small cetacean is heavily dependent on a continuous food supply (Hoekendijk et al., 2017; Wisniewska et al., 2016). This species is thus expected to be particularly sensitive to climate changes (Heide-Jørgensen et al., 2011). Besides, due their coastal distribution, thousands of harbor porpoises are also killed accidentally by commercial fisheries threatening local populations to a level that is still hard to quantify, but which is likely unsustainable especially with a highly fluctuating environment (Carlén et al., 2021; IMR-NAMMCO, 2019; Vingada & Eira, 2018).

Harbor porpoises are officially subdivided into three distinct subspecies: *P. p. vomerina* in the Pacific, *P. p. phocoena* in the North Atlantic, and *P. p. relicta* in the Black Sea (Fontaine, 2016; Read, 1999; Rosel et al., 1995). A fourth lineage, the Iberian-Mauritanian lineage (IBMA; possibly the unnamed sub-species *P. p. meridionalis*) has been suggested owing to their distinctive ecology, morphology (Donovan & Bjørge, 1995; Smeenk et al., 1992), and genetic divergence (Fontaine et al., 2014), but no formal description has been made for this distinct lineage (Pierce et al., 2020). IBMA is found around the relatively cold and productive upwelling waters off the coasts of Iberia and Mauritania. Iberian and Mauritanian porpoises are closely related but genetically distinct from each other and form two populations separated by at least 2500 km (Ben Chehida et al., 2021; Fontaine, 2016; Fontaine et al., 2014).

The upwelling habitat of the Iberian porpoise is at the border of a biogeographical transition zone between temperate and subtropical waters where species with different affinities co-occur and reach their respective ecological limits (Bowen, 1997; MacLeod, 2009). Because conditions at the margins between biogeographic zones are often suboptimal, populations like the Iberian harbor porpoises are expected to be among the first to show the impacts of such environmental changes. This is particularly true with the small scale Iberian upwelling system, which is known to fluctuate due to the ongoing climate change (Casabella et al., 2014; Pires et al., 2013). Such environmental fluctuation may impact the Iberian harbor porpoises in various ways, possibly leading to dramatic impacts on the population demography and genetic diversity. Porpoises may adapt locally for example by adapting to various prey species (Heide-Jørgensen et al., 2011), by migrating to more suitable environments, or the population may also become extirpated if environmental changes outpace their ability to adapt. Previous population genetic studies (Ben Chehida et al., 2021; Fontaine et al., 2014; Fontaine et al., 2010) showed that the Iberian population was a source population sending migrants Northward and Southward to neighboring populations, but received none. This observation was based on a limited sample size focusing on a cohort obtained about thirty years ago, between 1990 and 2002. If the recruitment does not keep up with the mortality and migration rate, the census size of the Iberian population is expected to decrease, leading to a gradual reduction in genetic diversity over time.

In this study, we investigated how the genetic diversity has changed in the Iberian harbor porpoises over the past three decades. More precisely, using genetic diversity as a proxy of population size variation, we tested if their genetic diversity decreased through time by comparing the genetic composition of two cohorts of porpoises, one sampled in the 1990s (1990-2002), and the other one from the 2010s (2012-2015).

## 2. Material and methods

### Sampling and molecular analyses

Tissue samples from 52 by-caught or stranded individuals between 2012 and 2015 were obtained from the Portuguese Marine Animal Tissue Bank (MATB) and were stored in 70% ethanol and kept at −20°C until molecular analyses. Genomic DNA was extracted with the DNeasy Blood and Tissue kit (QIAGEN Inc.) following the manufacturer’s protocol. Following Fontaine et al. (2014), we screened the mitochondrial genetic polymorphism for five coding regions (*ATP-6, ATP-8, Cyt-b, ND5*, and *COI*) using Sanger sequencing. Target regions were PCR amplified, quality controlled, and purified following the protocol described in Fontaine et al. (2014). Sanger sequencing was outsourced to GATC Biotech ltd. The resulting forward and reverse sequences were visually inspected, cleaned, and assembled with *Geneious* 10.0.9 (Kearse et al., 2012). We added 83 previously published sequences from Fontaine et al. (2014) encompassing the same five mitochondrial genes obtained for 82 harbor porpoises collected between 1990 and 2002 (Figure 1), and one sequence from the closest outgroup species, the Dall’s porpoise (*Phocoenoides dalli*). All mtDNA sequences were aligned with *MUSCLE* (Edgar, 2004). We grouped harbor porpoise sequences into five geographical regions (Figure 1): North Atlantic (n=23), Bay of Biscay (n=14), Iberia (n=71), Mauritania (n=14) and Black Sea (n=12). We further subdivided the Iberian porpoises into two cohorts: new (n=52, 2012-2015) vs. old (n=19, 1990-2002). The temporal coverage of the sampling is thus at most 25 years, which corresponds to 2.5 generations, assuming a generation time of 10 years (Read, 1999).

**Figure 1.**
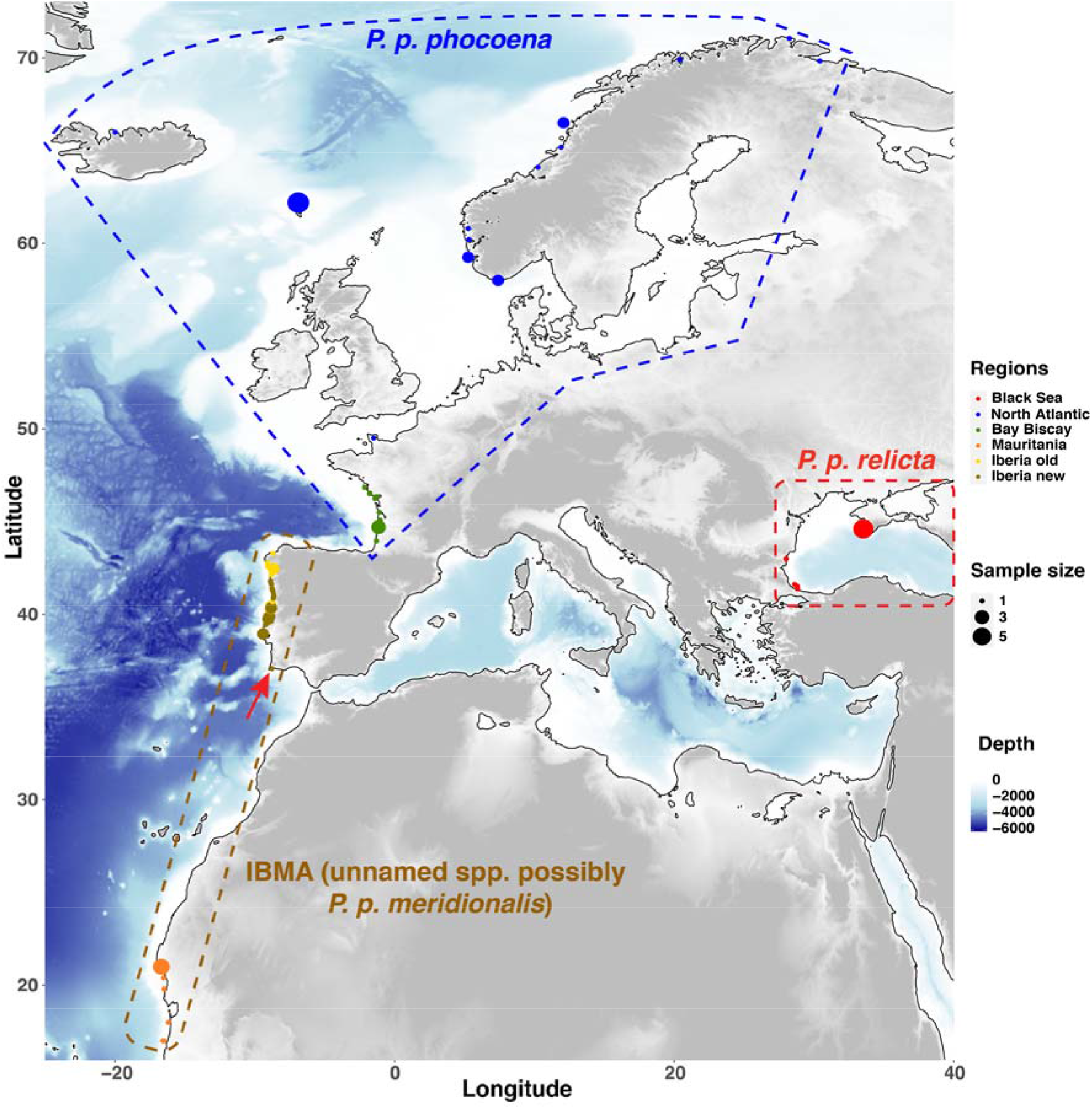
Map showing the sampling locations and sample sizes. Geographic locations are based on approximate GPS coordinates or reported discovery locations. The red arrow indicates the geographical location of the two porpoises carrying Haplotype 10.

### Phylogenetic and population genetic analyses

In order to place the Iberian porpoises into a global phylogeographic context, we first reconstructed a maximum-likelihood (ML) phylogeny with all the 135 mtDNA sequences. Unique mtDNA haplotypes were identified using *DnaSP* v.5.10.1 (Librado & Rozas, 2009). Then, we used *PhyML* (Guindon et al., 2010) implemented in *Geneious* to estimate the ML tree based on all available unique haplotypes identified in the dataset using a nearest neighbor interchange typology search. The support of each node was assessed using 5000 bootstrap resampling. The phylogenetic tree was rooted with the sequence from a Dall’s porpoise. In addition to the phylogenetic trees, we also reconstructed a median-joining haplotype network (Bandelt et al. 1999) using *PopART* (http://popart.otago.ac.nz).

Various statistics informative on the genetic variation of the two cohorts of Iberian porpoises (old vs. new) were estimated using the C library *libdiversity* (https://bioinfo.mnhn.fr/abi/people/achaz/cgi-bin/neutralitytst.c). Genetic diversity was estimated using the number of haplotypes (*H*), nucleotide diversity (*π*), and Watterson’s theta (*θ*_*W*_). A rarefaction approach was applied to estimate measures of genetic diversity while correcting for difference in sample size (Sanders, 1968) using a custom python script. Subsequently, genetic differentiation between the two cohorts was assessed using the Hudson’s estimator of *F*_*ST*_ (Hudson et al., 1992) and *φ*_*ST*_ (Excoffier et al., 1992) using respectively *DnaSP* v.5.10.1 (Librado & Rozas, 2009) and *Arlequin* v.3.5.2.2 (Excoffier & Lischer, 2010). The significance levels of *F*_*ST*_ and *φ*_*ST*_ were assessed by randomization tests (1,000 permutations). Finally, we investigated evidence of changes in mtDNA effective population size (*Ne*) between the two cohorts (old vs. new) by calculating values of Tajima’s *D* (Tajima, 1989) and Achaz’s *Y* (Achaz, 2008) by implementing also the rarefaction approach accounting for differences in sample sizes.

## 3. Results and Discussion

In total, we successfully sequenced 52 Iberian harbor porpoises from the new cohort (2012-2015). When aligned with the 83 sequences from Fontaine et al. (2014), the final alignment of 4,175 base-pairs (pb) included 135 individuals and contained 384 segregating sites with 267 singletons and 117 parsimony informative sites defining 61 distinct haplotypes (Table S1).

Consistent with previous studies, the phylogenetic relationships among mtDNA haplotypes depicted by the ML tree (Figure 2) and the haplotype network (Figure S1) revealed the three main monophyletic lineages corresponding to previously identified distinct subspecies (Fontaine et al., 2014): (i) *P. p. relicta* in the Black Sea; (ii) *P. p. phocoena* composed of the porpoises from the North Sea to the waters of Norway; and (iii) IBMA (unnamed *ssp*. possibly *P. p. meridionalis*), which comprises two distinct monophyletic sub-lineages in the upwelling waters of Iberia and Mauritania.

**Figure 2:**
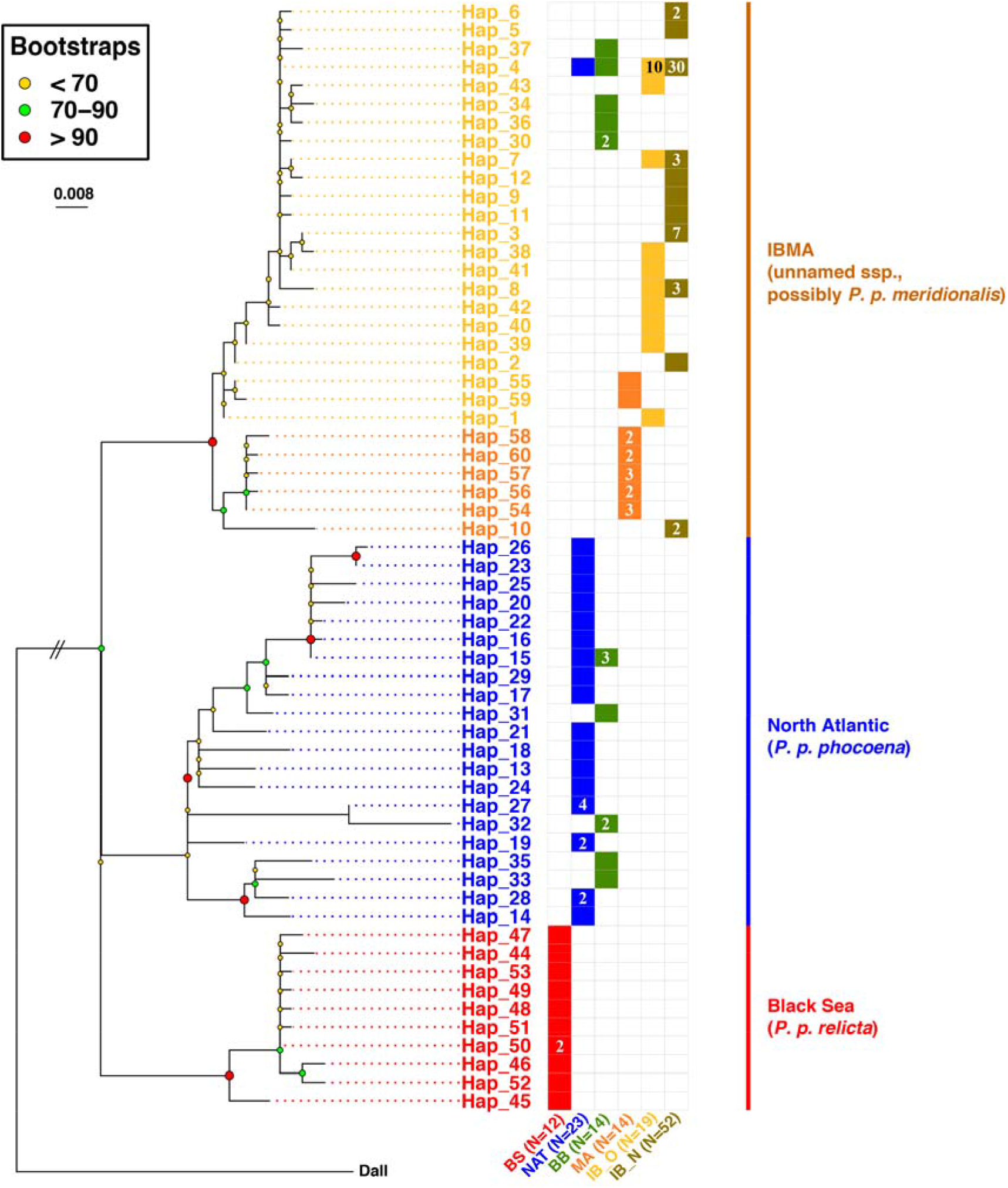
Maximum-likelihood mitochondrial phylogeny among unique mitochondrial haplotypes. Poorly supported nodes with less than 60% bootstrap support were collapsed. The color-coded labels show the geographic origin of the haplotype. The numbers within the boxes refer to the number of individuals carrying this haplotype. No number in a box means that the haplotype was observed only once. BS=Black Sea; NAT=North Atlantic; BB=Bay of Biscay; MA=Mauritania; IB_O=Iberian Old; IB_N= Iberian new.

Evidence of migration from Iberia to Mauritania has been previously documented (Ben Chehida et al., 2021; Fontaine et al., 2014) and is also observed here (Figure 2 and S1). However, we also identified one haplotype (Haplotype 10, see Figure 2 and S1) geographically present in the new Iberian cohort, but clustering with the Mauritanian lineage. This haplotype was carried by two individuals in the southernmost part of the Iberian sampling (red arrow in Figure 1). This result provides the first evidence of possible migration from the Mauritanian population into Iberian waters, suggesting that these two populations may not be as isolated as previously thought. Haplotype 10 could also belong to a distinct unknown population as it is relatively divergent from the other Mauritanian haplotypes (Figure 2 and S1). Indeed, the ML tree (Figure 2) shows that this haplotype forms a distinct lineage from the other Mauritanian porpoises. The haplotype network also illustrates this divergence (Figure S1): while the Mauritanian haplotypes are separated from each other with at most two mutational steps, at least ten mutational steps separate this haplotype 10 from the other Mauritanian haplotypes (Figure S1). Notably, both Iberian porpoises carrying this haplotype 10 were found on the southern extremity of the Portuguese coasts (8.78W-37.07N; Praia do Burgau, Portugal), at least 400km south from all other Iberian sampling locations (Figure 1). Stranding records of harbor porpoises were previously reported in Morocco, on the east coasts of Gibraltar, and in Cadiz (Spain; Rojo–Nieto et al., 2011). If unsampled populations occur between the Iberian Peninsula and the Mauritanian waters, they are most likely confined to a narrow and seasonal upwelling area. This relatively divergent haplotype 10 closely related to the Mauritanian rather than the Iberian porpoises could be evidence of these unsampled populations. It is all the more possible that harbor porpoise sightings, stranding, and bycatches have been continuously reported from the North of Morocco to Gambia (Boisseau et al., 2010; IMR-NAMMCO, 2019). Haplotype 10 was excluded from subsequent analyses of genetic diversity because it could be part of a distinct population and therefore would artificially inflate measures of genetic diversity (Table S1). Future studies with additional samples and data from the nuclear genome are needed to confirm whether these two porpoises in the southernmost part of Iberia are genetically distinct from the other Iberian populations, possibly forming a distinct population, and whether or not they are admixing with the other Iberian porpoises.

Aside from the two porpoises carrying the haplotype 10, our results indicate that both cohorts sampled at different times along the Iberian coasts belong to a same mtDNA genetic pool. We detected no genetic differentiation suggesting no significant differences in haplotype frequencies between them (*F*_*ST*_ and *φ*_*ST*_ values < 0.0025; *p*-value > 0.08). This result reflects the high prevalence of haplotype 4 in both cohorts (Figure 2 and S1). However, we observed a dramatic reduction in genetic diversity in the new cohort compared to the old one as shown by the lower number of haplotypes (*H*; Figure 3a), and in the two measures of nucleotide diversity (*π* and *θ*_*W*_; Figure 3b and 3c, respectively). The rarefaction procedure (to account for differences in sample size between the two cohorts) provided striking evidence of this reduction (Figure 3). The non-overlapping standard errors from a standardized sample size equal to (or larger than) eight sequences in the rarefaction curves suggest that the decline of genetic variation is highly significant. For example, the average number of observed haplotypes (*H*) observed with 19 randomly sampled sequences in each cohort decreases from 10 to 6 when comparing the old vs. new cohorts. This suggests that the proportion of rare haplotypes has dropped dramatically over the 25-years sampling period of this study. Consistent with this observation is the rapid loss in nucleotide genetic diversity as measured by *π* and *θ*_*W*_ (Figure 3b and 3c). A stochastic loss of genetic variation of this magnitude is often attributed to an enhanced effect of genetic drift (Hartl, 2020), which could suggest a dramatic population decline.

**Figure 3:**
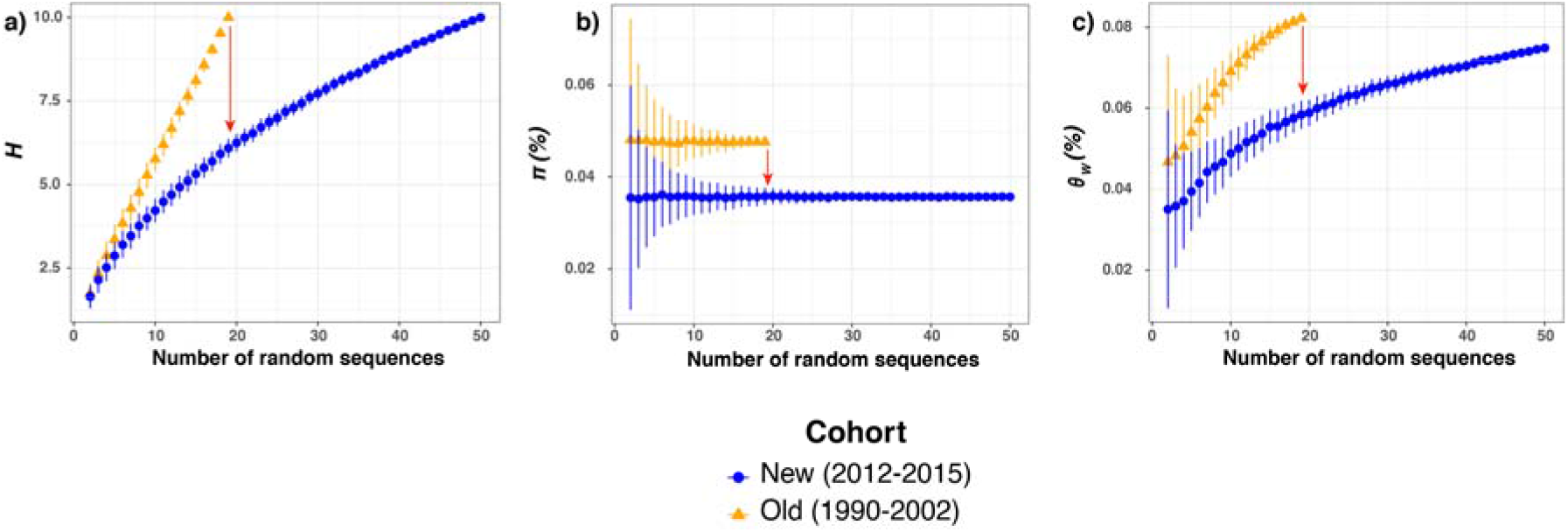
Rarefaction curves showing three statistics describing mitochondrial genetic diversity including (a) number of haplotypes (*H)*, (b) the nucleotide diversity (*π*), and (c) Watterson’s theta (*θ*_*W*_). The red arrow in each panel shows the direction of change between the old and new cohorts. The X-axis shows the number of sequences randomly resampled in the rarefaction procedure to estimate the statistical values.

Tajima’s *D* (Tajima, 1989) and Achaz’s *Y* (Achaz, 2008) statistics estimated for the two cohorts sampled at different times along the Iberian coast present negative values (Figure 4). At first glance, such negative values would be indicative of a population that underwent a “recent” expansion at evolutionary time-scales (Epps & Keyghobadi, 2015; Smith et al., 2011), most likely due to the foundation of the Iberian population during the period that followed the Last Glacial Maximum (19kyr ago; Fontaine et al. 2014; Fontaine 2016). However, the values estimated for the new cohort for both statistics were significantly less negative than those from the old cohort. For example, the Tajima’s *D* values obtained with 19 randomly sampled sequences increased from −1.5 for the old cohort to −1.25 for the new (Figure 4a). Likewise, values of Achaz’s *Y* statistic, which account for possible sequencing errors (Achaz, 2008), show an even stronger increase from −1.75 to −0.95 between the old and new cohorts (Figure 4b). It is remarkable to see how both values have become dramatically less negative in the time lapse of only 2.5 generations between the two cohorts. This increase in *D* and *Y* values also provides a strong indication of a population losing rapidly rare haplotypes as expected for a small and isolated population strongly subject to genetic drift that could be declining.

**Figure 4:**
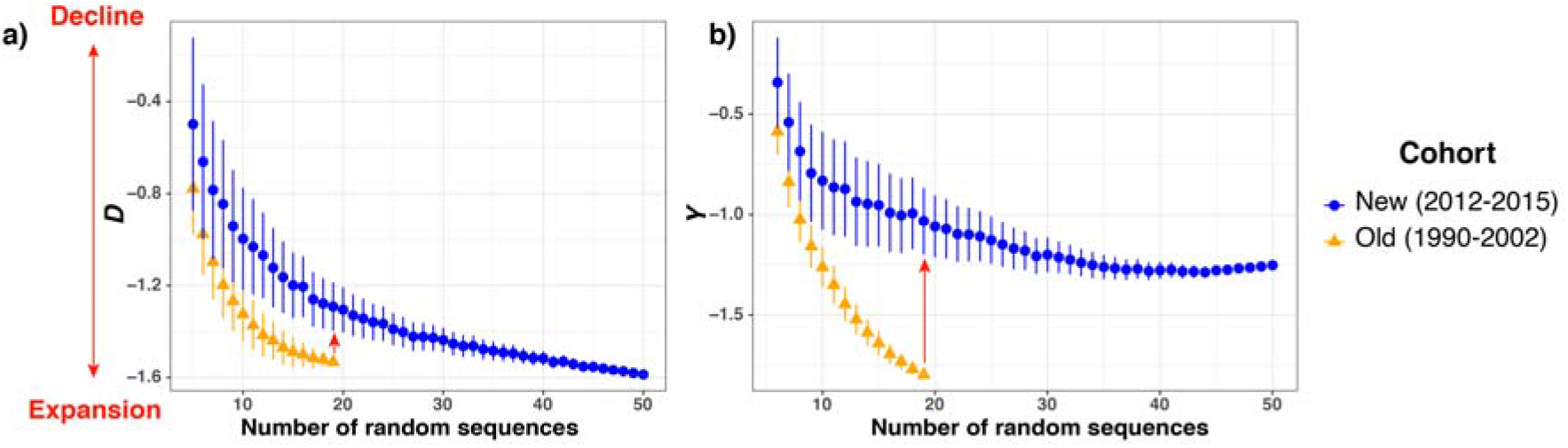
Rarefaction curves showing two neutrality indexes indicative of demographic changes including (a) Tajima’s *D* and (b) Achaz’s *Y*. The red arrow inside each panel shows the direction of the change between the old and new cohorts. The bidirectional arrow shows the interpretation of *D* and *Y* in terms of demography. The X-axis shows the number of sequences randomly resampled in the rarefaction procedure to estimate the statistical values.

We confirmed here previous evidence of gene flow from Iberia to more northern regions with Iberian haplotypes present in the Bay of Biscay and in the English Channel where admixture between the two lineages, IBMA (or *P. p. meridionalis*) and *P. p. phocoena*, have been reported (Figure 2 and S1; Ben Chehida et al., 2021; Fontaine et al., 2014; Fontaine et al., 2017). Additionally, like previous studies (Ben Chehida et al., 2021; Fontaine et al., 2007; Fontaine et al., 2014; Fontaine et al., 2017; Fontaine et al., 2010), we did not find evidence of gene flow from the population located further North into IBMA. Because the population size of *P. p. phocoena* in the North Sea is much larger than that of the Iberian population, the observed northward migration may lead to the assimilation of the Iberian gene pool by *P. p. phocoena*. In the long term, this process may lead to the disappearance of the unique Iberian genetic pool. This scenario is supported by Macleod et al. (2009), which used habitat modeling to show that species with a similar range as the Iberian harbor porpoises (Figure 1C in MacLeod, 2009) may shift their range northward as a response to an increase in water temperature of 5°C due to global warming (Figure 1D in MacLeod, 2009). This model suggests that the future range of IBMA (*P. p. meridionalis*) may totally overlap with the current range of *P. p. phocoena* (Figure 1D in MacLeod, 2009). It is worth mentioning that Ben Chehida et al. (2021) recently modeled the future distribution of the harbor porpoises in the North Atlantic under the most aggressive scenario of the Intergovernmental Panel on Climate Change (IPCC) in 2050 (RCP8.5; Schwalm et al., 2020) and did not capture this strong northward range shift. A plausible explanation of the discrepancy between the two studies is related on one side to the temperature increase assumed by the two models, 2°C and 5°C, in Ben Chehida et al. (2021) and MacLeod (2009), respectively. On the other side, the environmental niche modeling in Ben Chehida et al. (2021) did not account for any biotic factors in the simulations.

Hence, all the results of this study support the hypothesis that the Iberian population may be small and isolated. Furthermore, both the sharp decline in genetic diversity over the past 25 years and northward migration can represent sources of concern for the long-term survival of the Iberian porpoises. On one hand (hypothesis 1), the decay of genetic variation we reported here could simply reflect the fact that Iberian porpoises constitute a stable population with a very low effective population size (*Ne*). In such a condition, genetic drift has a disproportionate influence on the genetic make-up of this population leading to the observed loss of genetic variation over a few generations. Several previous studies showed that the Iberian population has indeed a very low *Ne* at nuclear microsatellites loci (Ne < 100 individuals; Ben Chehida et al., 2021; Fontaine et al., 2014; Fontaine et al., 2010). The mtDNA locus employed in this study has an effective size four times smaller compared with nuclear autosomal loci. This means that the stochastic loss of genetic variation due to genetic drift is expected to be drastically increased for the mtDNA compared with the nuclear loci. Thus, the Iberian population might be at a stable census size losing rare haplotypes through the enhanced effect of genetic drift at the mtDNA locus. On the other hand (hypothesis 2), the decay of genetic variation might be associated with a rapid decline of the number of reproductive females in addition to the overall low *Ne* (Lande, 1988). This hypothesis is even more plausible since there is accumulating evidence that the Iberian population is sending more migrants to adjacent populations than the reverse. This second hypothesis is concerning because, in addition to being small and isolated, the decline of the census population would imply that the Iberian porpoises are becoming increasingly subject to demographic stochasticity (Allendorf et al., 2012). Because this population is also known to be subject to strong genetic stochasticity (i.e., genetic drift; see this study) and environmental stochasticity (Casabella et al., 2014; Pires et al., 2013), all these effects can accumulate and reinforce each other, potentially precipitating the decline of the Iberian porpoises. This so-called extinction vortex (Allendorf, et al., 2012) can be particularly severe for a long-lived *K* strategist species, inhabiting a marginal habitat highly subject to environmental variation like the Iberian harbor porpoises (Casabella et al., 2014; Pires et al., 2013) and could put them at a high risk of extinction.

This study relies on a single molecular marker (mtDNA). Therefore, future studies with additional samples and data from the nuclear genome are highly needed to tease apart the two hypotheses depicted above. Indeed, genome-wide data can provide further insights into the temporal dynamics of the Iberian population. For example, the reconstruction of historical demography based on genome-wide data (Liu & Fu, 2020; Schiffels & Wang, 2020; Terhorst et al., 2017) can be used to test whether these harbor porpoises, since their post-glacial emergence (Fontaine, 2016; Fontaine et al., 2014; Fontaine et al., 2010), maintained a long-term low *Ne* (Morin et al., 2020; Robinson et al., 2019) or if, on the contrary, they declined recently (Hu et al., 2020). A simulation framework, similar to the one developed by Robinson et al. (2019), could be used to assess whether these porpoises managed to escape deleterious effects of the mutation load despite their current low *Ne* or not, as recently suggested for the vaquita (*P. sinus*; Morin et al., 2020; Robinson et al., 2019).

Additional sources of concern possibly related to a severe decline in the Iberian population require careful considerations: the SCANS III project estimated a census size as low as 2,898 animals (CV=0.32) in Iberian waters (Hammond et al., 2017). Also, census estimates for Portuguese waters for a continuous 6-year time series (2010-2015) led to an estimated abundance of only 1,531 individuals (CV=0.31) (Vingada & Eira 2018) with concerning interannual fluctuations. These low census sizes were associated with a particularly high bycatches rates with best estimates varying from 90-197 deaths per year (Pierce et al., 2020) to 130-330 deaths per year (Read et al., 2020). However, it is important to mention that these estimates are mainly based on on-board observer data that have very wide confidence limits due to low observer coverage of fishing activities in the region and all estimates (from observer data, strandings, and interviews) are potentially biased. There may also be overexploitation of feeding resources (Méndez-Fernandez et al., 2013; Santos & Pierce, 2003). A scarcity in prey availability can be harmful because it enhances interspecific competition, as already reported between harbor porpoises and other marine mammals, for example with the bottlenose dolphin *Tursiops truncatus* which ecological niche largely overlaps with the harbor porpoise in some portions of the Iberian waters (Méndez-Fernandez et al., 2013; Spitz et al., 2006). Hostile interactions between the two species have been documented in Iberian waters and hypothesized to cause harbor porpoises to disperse to less productive narrow shelf areas (Alonso et al., 2000; Spitz et al., 2006). Furthermore, Iberian harbor porpoises are regularly found stranded, often with signs of bycatch mortality, potentially indicating unsustainable mortality rates (IMR-NAMMCO, 2019; López et al., 2002; Pierce et al., 2020; Read et al., 2020; Vingada & Eira, 2018). In fact, up to 15% of the harbor porpoise population may be taken annually by fisheries (particularly gill and trammel nets) operating in Portuguese waters (Vingada & Eira, 2018).

While the IBMA lineage is known to be unique in terms of genetics, morphology, and ecology, they are currently not recognized as a distinct subspecies (Pierce et al., 2020). There is also no mention of these porpoises in the IUCN Red List of Threatened Species or the List of Marine Mammal Species and Subspecies of the Society of the Marine Mammalogy despite the multiple threats faced by this vulnerable lineage. Given the results of this study and the previous concerns raised about their viability (IMR-NAMMCO, 2019; Pierce et al., 2020), we recommend immediate actions from the community to officially recognize these porpoises as a distinct subspecies and to receive a formal conservation assessment from the IUCN. Furthermore, it is paramount to ensure that all the measures are taken to gain additional knowledge and to apply effective conservation efforts to protect the Iberian population. Recently, Spain changed the conservation status of the harbour porpoises inhabiting Spanish waters from Vulnerable to Endangered (Boletin Oficial del Estado, 2020). In Portugal, a new Marine Protected Area (PTCON0063) in the northwestern coast was recently designated whereas another one was enlarged in the southwestern coast (PTCON00012) in order to incur higher protection to harbor porpoise hotspots in the country (Vingada & Eira, 2018). Nonetheless, none of the conservation and mitigation measures within the areas’ management plan (Portaria 201/2019) is currently in place and bycatch mortality remains effectively unaddressed. In the future, improving our understanding of the demographic trends and the threats these porpoises are facing is paramount to devise suitable and tailored larger scale conservation plans for the whole Iberian population. Such measures are timely needed to mitigate the decline of this unique lineage potentially on the brink of extinction.

## Acknowledgements

We thank all the people and networks involved in the sampling collection of harbor porpoises over the years, including KA Tolley, T Jauniaux, J Haelters, AA Llavona, B and AA Ozturk, V Ridoux, F Caurant, W Dabin, E Rogan, M Sequeira, U Siebert, GA Vikingsson, A Borrell, AA Aguilar, A Marçalo, and the national stranding networks: Réseau National d’Échouage and PELAGIS in France; Royal Belgian Institute of Natural Sciences-MUMM, and Dept. Pathology of the U. Liège, Belgium; CEMMA-Coordinadora para o Estudio dos Mamiferos Mariños, Spain; the Portuguese Marine Animal Tissue Bank-MTAB, Portugal; the Dutch Stranding Network, SOS Dolfijn, and Dept. Pathology of the U. Utrecht in the Netherlands; and the Dept. pathology of the U. Kiel, Germany. All samples were collected under appropriate permits following the relevant guidelines and regulations, and transferred internationally under CITES permits. We also thank Graham J. Pierce for his feedbacks on the first draft of the manuscript and the participants of the IMR-NAMMCO workshop in Tromsø (Norway, December 2018) for their significant inputs during the development of this study. We are also grateful to the Centre for Information Technology of the University of Groningen, and in particular Bob Dröge and Fokke Dijkstra, for their continuous support and for providing access to the Peregrine high-performance computing cluster. This study was funded by the University of Groningen (The Netherlands). YBC was supported by a PhD fellowship from the University of Groningen.

## Data Availability

The data sets and scripts supporting this study are available in IRD Porpoises Genetics and Genomics Dataverse Repository (https://dataverse.ird.fr/dataverse/porpoise_genomics) at doi: To Be Annouced (TBA). Mitochondrial haplotypes were also deposited on the NCBI– GenBank under the Accession Numbers (Accession numbers : TBA).

## Author Contributions

Conceptualization: MCF. Data curation: MF, ATP, LN, MCF. Formal analysis: YBC, TS, MCF. Funding acquisition: MCF. Investigation: JT, TS, JPAH, MCF. Methodology: JT, TS, JPAH, MCF. Project administration: MCF. Resources: JT, TS, JPAH, MCF. Software: YBC. Supervision: MCF. Validation: MCF, YBC, JT. Visualization: TS, YBC, MCF. Writing – original draft: YBC, TS, MCF. Writing – review & editing: YBC, MCF with inputs from all the co-authors.

