## Supplementary Information for "Harbor porpoise losing its edges: genetic time series suggests a rapid population decline in Iberian waters over the last 30 years"

**ELECTRONIC SUPPLEMENTARY MATERIAL**

**Table**

**Table S1**: Mitochondrial genetic diversity

|  | **All  (with outgroup)** | **All  (without outgroup)** | **BS** | **NAT** | **BB** | **MA** | **All IB** | **IB_O** | **IB_N** | **IB_N (excl. hap 10)** |
| --- | --- | --- | --- | --- | --- | --- | --- | --- | --- | --- |
| *N* | 135 | 134 | 12 | 23 | 14 | 14 | 71 | 19 | 52 | 50 |
| *S* | 384 | 170 | 23 | 99 | 66 | 11 | 27 | 12 | 24 | 14 |
| *Singl.* | 267 | 65 | 21 | 57 | 19 | 1 | 7 | 6 | 5 | 7 |
| *Parsim.* | 117 | 105 | 2 | 42 | 47 | 10 | 20 | 6 | 19 | 7 |
| *# Hap* | 61 | 60 | 10 | 18 | 10 | 7 | 17 | 10 | 11 | 10 |
| *π (%)* | 0.50 | 0.40 | 0.10 | 0.41 | 0.52 | 0.07 | 0.05 | 0.05 | 0.06 | 0.04 |
| *θ_W_ (%)* | 1.68 | 0.74 | 0.18 | 0.64 | 0.50 | 0.08 | 0.13 | 0.08 | 0.13 | 0.08 |
| *D* | ~~-~~ | - | -2.04 | -1.46 | 0.22 | -0.66 | -1.85 | -1.53 | -1.77 | -1.58 |
| *Y* | ~~-~~ | - | -1.56 | -0.83 | 0.05 | -1.93 | -2.12 | -1.77 | -2.19 | -1.26 |

*N*, mtDNA sample sizes; *S*, segregating sites; *Singl*., singleton; *Parsim*., sites informative in parsimony; *# Hap*, number of haplotypes; *π*, nucleotide diversity; *θ_W_*, Watterson's theta; *D*, Tajima's *D*; *Y*, Achaz's *Y*. BS=Black Sea; NAT=North Atlantic; BB=Bay of Biscay; MA=Mauritania; IB_O=Iberian Old; IB_N= Iberian new.

**Figures**

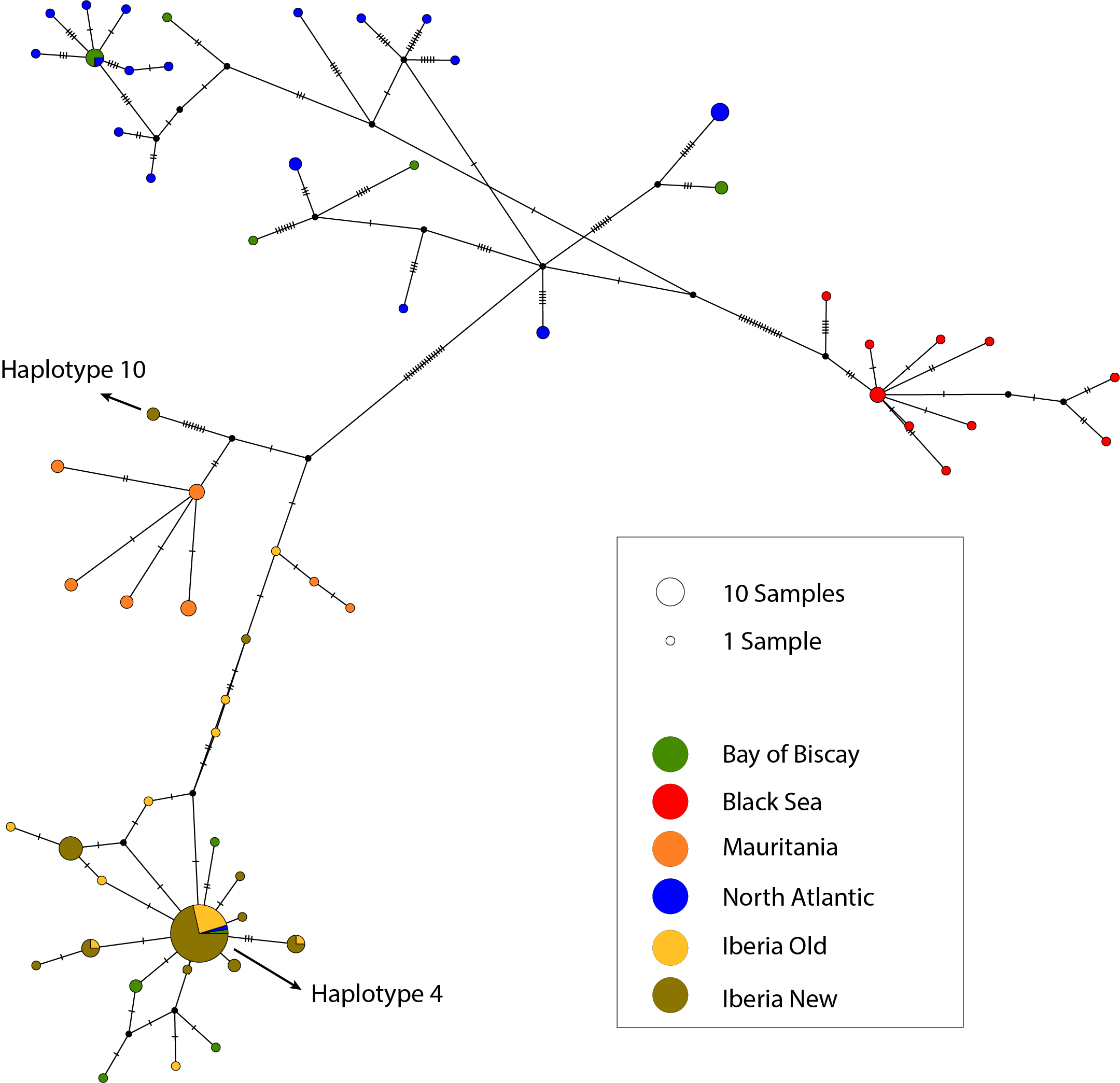

**Figure S1.** Mitochondrial median-joining haplotype network. Each circle represents a haplotype. The size of each circle is proportional to the haplotype frequency observed in the total sampling, and each pie slice is proportional to the number of each haplotype observed per geographic location. Each dash represents a unique mutational step between haplotypes. A black dot represents a hypothetically unsampled or extinct node. The arrows highlight haplotypes 4 and 10 discussed in the main text.
